# An active finite viscoelastic model for gastric smooth muscle contraction

**DOI:** 10.1101/2021.01.26.428273

**Authors:** Satish Kumar Panda, Martin Lindsay Buist

## Abstract

A coupled electromechanical model to describe the transduction process of cellular electrical activity into mechanical deformation has been presented. The model consolidates a biophysical smooth muscle cell model, a biophysical actin-myosin interaction model, a sliding filament model and a viscoelastic constitutive model to construct an active finite viscoelastic model. The key input to this model is an electrical pulse which then estimates the resulting stress and deformation in the cell. The proposed model was used to recreate experimental observations performed on canine and porcine gastric tissue strips. In all cases, the simulation results were well matched with the experimental data (*R*^2^ *>* 0.9).

## 1 Introduction

Smooth muscle cells (SMC) are the effectors of motility in gastrointestinal (GI) tract. In an SMC, the contractile units (CU) are the force-producing elements, and they consist of a thick myosin filament surrounded by thin actin filaments. An elevation in intracellular *Ca*^2+^ concentration ([*Ca*^2+^]_*i*_) activates the enzyme myosin light chain kinase which in turn triggers phosphorylation and conformational changes in the myosin light chain. This process forms cross-bridges between actin and myosin filaments. Different models have been proposed to demonstrate this process mathematically [5, 7, 13]. The newly formed cross-bridges force the actin and myosin filaments to slide relative to each other and initiate contraction in the cell. The force produced in this process is commonly referred as the active force that is often quantified using sliding filament models [9, 15, 26]. Therefore, for a quantitative description of how the cellular activities transduce into a mechanical deformation, it is necessary to construct an electromechanical framework.

In recent years a small number of simulation studies have appeared in the literature to describe the electromechanics of the GI tract [3, 4, 15]. The work by Kroon, for example, quantified the active force produced by the GI SMC when stimulated by an external *Ca*^2+^ signal [9, 10]. He consolidated a viscoelastic model with the four-state model of Hai and Murphy to examine the transduction pathway from *Ca*^2+^ signal to mechanical deformation. With this model, the author was able to recreate the uniaxial stress-stretch behaviour of guinea pig taenia coli strips under isometric and isotonic loading conditions. However, the author did not consider the sliding filament theory in his work. Also, the four-state model of Hai and Murphy is one of the earliest models for actin-myosin interaction, and in its original form, it may not be suitable for describing the cross-bridge formation process in the GI SMC. Furthermore, the author utilised empirical relations to relate the external *Ca*^2+^ signal to the intracellular *Ca*^2+^ concentration variation inside the SMC which may not describe the true behaviour of the cell. In this context, Murtada et al. proposed a method to compute the active forces while considering the sliding filament theory; however, they did not incorporate any cellular electro-physiology model or cellular contraction model in their work [15, 16]. The study by Du et al. was able to demonstrate the active force produced by GI SMC when stimulated by external electrical pulses; however, their work had been carried out with the contraction and material models originally proposed for cardiac electromechanics [4]. Also, the authors utilized a hyperelastic material for the GI tissue which may not provide the true tissue properties as the GI tissue properties are nonlinear and viscoelastic [20]. Hence, we propose a novel electromechanical model, here, to compute active forces in GI SMC by coupling a cellular electrophysiology model, an actin-myosin interaction model, a sliding filament theory, and a nonlinear hyper-viscoelastic model together.

## 2 Methods

We use the notations *a*, **a** and **A** for a scalar, a vector and a second-order tensor, respectively. The trace and the deviatoric part of a tensor are denoted as: tr(**A**) = **I** : **A, A**^*D*^ = **A** - (1/3) tr(**A**) **I**, where the (:) represents a tensor scalar product and **I** is the second order identity tensor. The deformation gradient for a continuum body can be expressed as **F** = *∂****x*** */∂****X*** where ***X*** and ***x*** are the material points in the continuum body in the reference (*Ω*_*o*_) and current configurations (*Ω*), respectively. Other important deformation tensors may be defined as: the right Cauchy-Green deformation tensor, **C**= **F**^*T*^ **F**; the left Cauchy-Green deformation tensor, **B**= **FF**^*T*^ ; and the velocity gradient tensor, 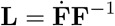, where the 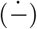 represents the material time derivative. The stress tensors (e.g., Cauchy stress tensor, **S**; first Piola-Kirchhoff (PK) stress tensor, **P**; and second PK stress tensor, **T**) are also related to each other through **F**, i.e., **S** = (1*/*det (**F**))**FTF**^*T*^, **T** = **PF**^*−T*^.

### 2.1 Actin-myosin interaction model

The actin-myosin interaction is the primary means to generate contractile activity in a SMC. The four-state model proposed by Hai and Murphy is the most popular model to explain the actin-myosin interaction in SMC [7]. However, a bimodular biophysical based model proposed by Gajendiran and Buist for gastric SMC was adopted in this work as it was able to explain the relationship between [*Ca*^2+^]_*i*_ and contraction in the canine SMC accurately [5]. Given an input [*Ca*^2+^]_*i*_, this model calculates the total number of cross-bridges in the SMC and also the contractile force generated inside the cell. However, this model assumes that the contractile force produced in the cell is proportional to the number of cross-bridges which clearly ignores sliding filament theory. Therefore, we need to couple it with a suitable sliding filament model.

### 2.2 Sliding filament model

Smooth muscle generates active force over a wide range of muscle stretch. The active force produced by a CU is stretch-dependent and the active force-length relation is bell-shaped. At an optimum stretch value, 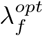, the active force becomes maximum and then it decreases in both directions from 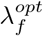 [15]. Fig. 5a shows an experimental observation of active force in a porcine taenia coil which clearly illustrates that the active force is maximum at a stretch of value 1.75 which is the optimum stretch for this tissue.

**Fig 1:**
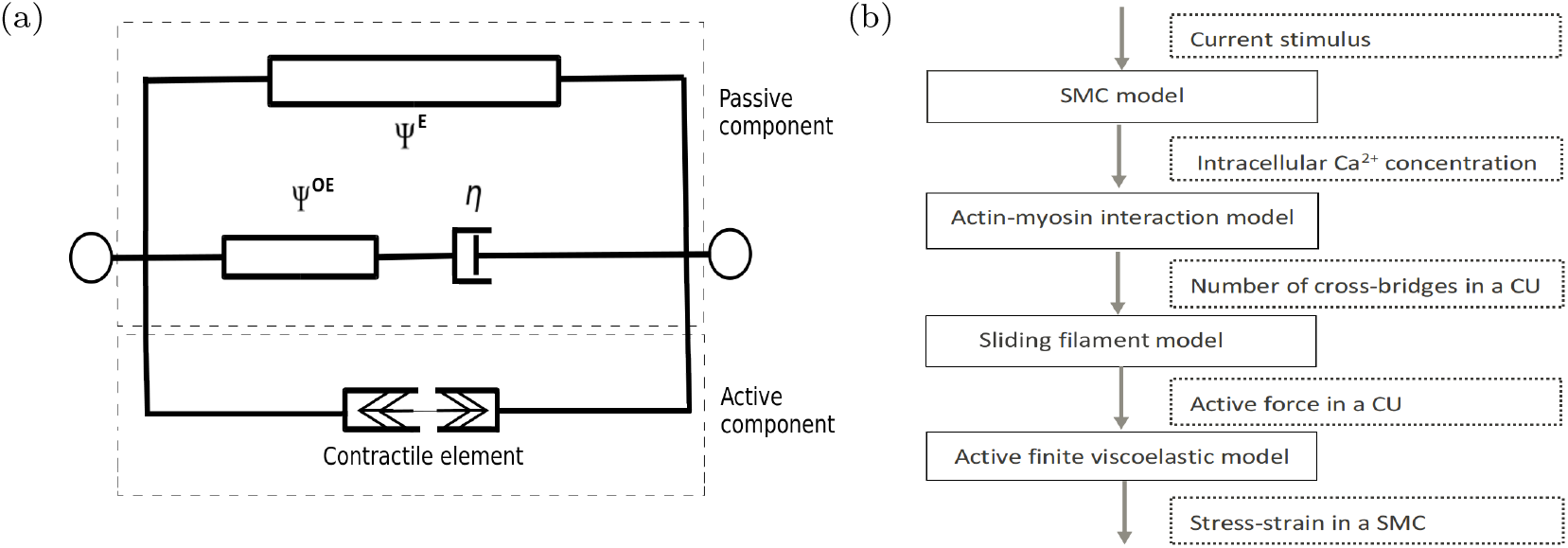
(a) A rheological model for the active finite viscoelastic model. In the figure, *ψ*^*E*^ and *ψ*^*OE*^ are the strain energies for the in-series and parallel hyperelastic elements, respectively, and *η* is the viscosity of the dashpot. (b) A schematic diagram showing the coupling of different models to form the AFNHVM. The entities in dotted square brackets are the input/output of a model.

**Fig 2:**
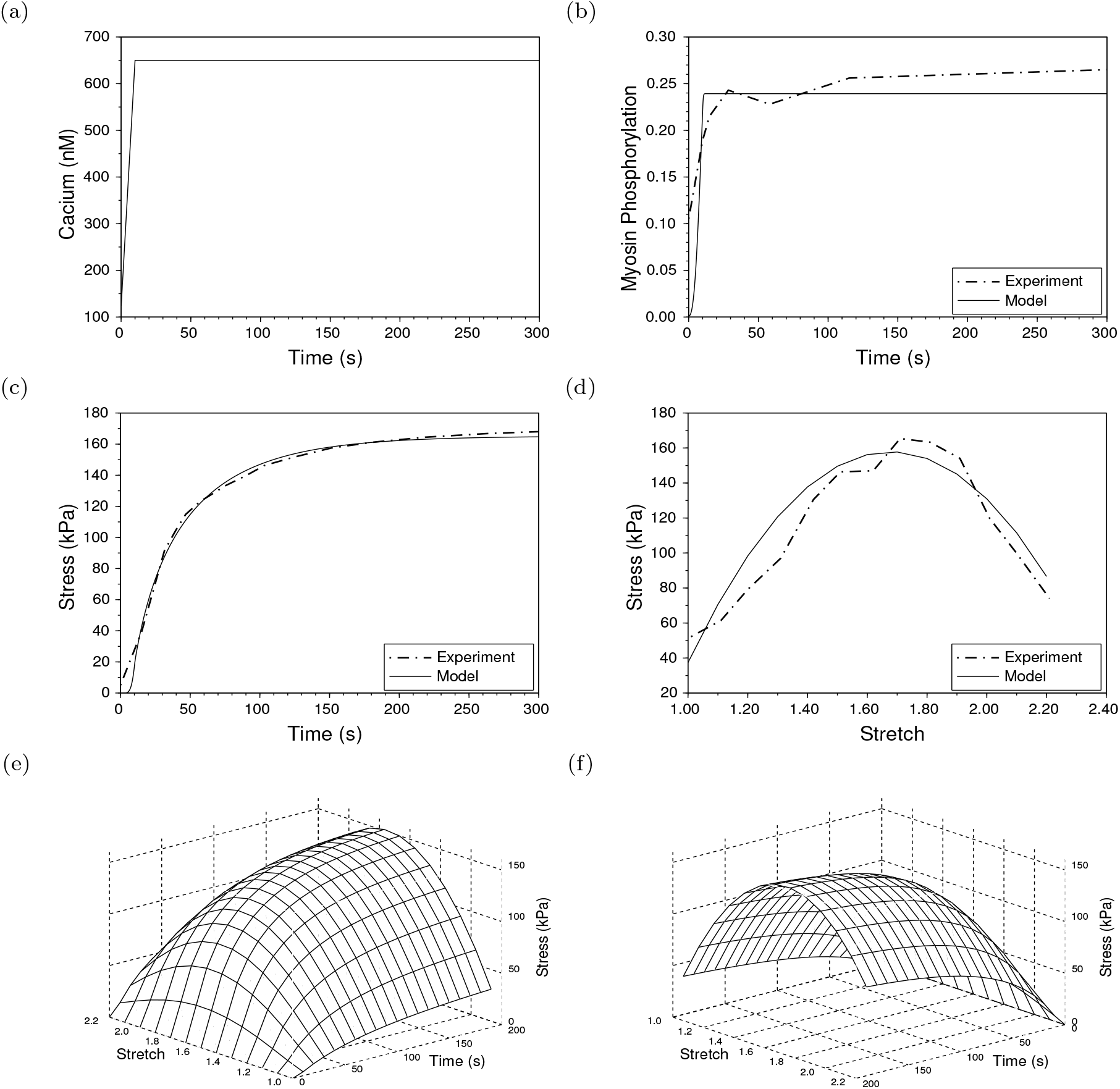
(a) A step change in [*Ca*^2+^]_*i*_ that served as an input to the AFNHVM model. (b) The experimental and model predicted myosin phosphorylation levels in the SMC [6]. (c) The experimental and model predicted evolution of active force with respect to time [6]. The optimized fitting parameters were *α* = 113.174 kPa, *β* = 0.0353 min^*−*1^ and *κ* = 2098.37 kPa. (d) Variation of active force with stretching lengths [24]. (e,f) The complete active response of a gastric tissue strip for different muscle lengths (*λ*_*f*_ = 1.0 *−* 2.2) from the time of activation up to 200 seconds.

**Fig 3:**
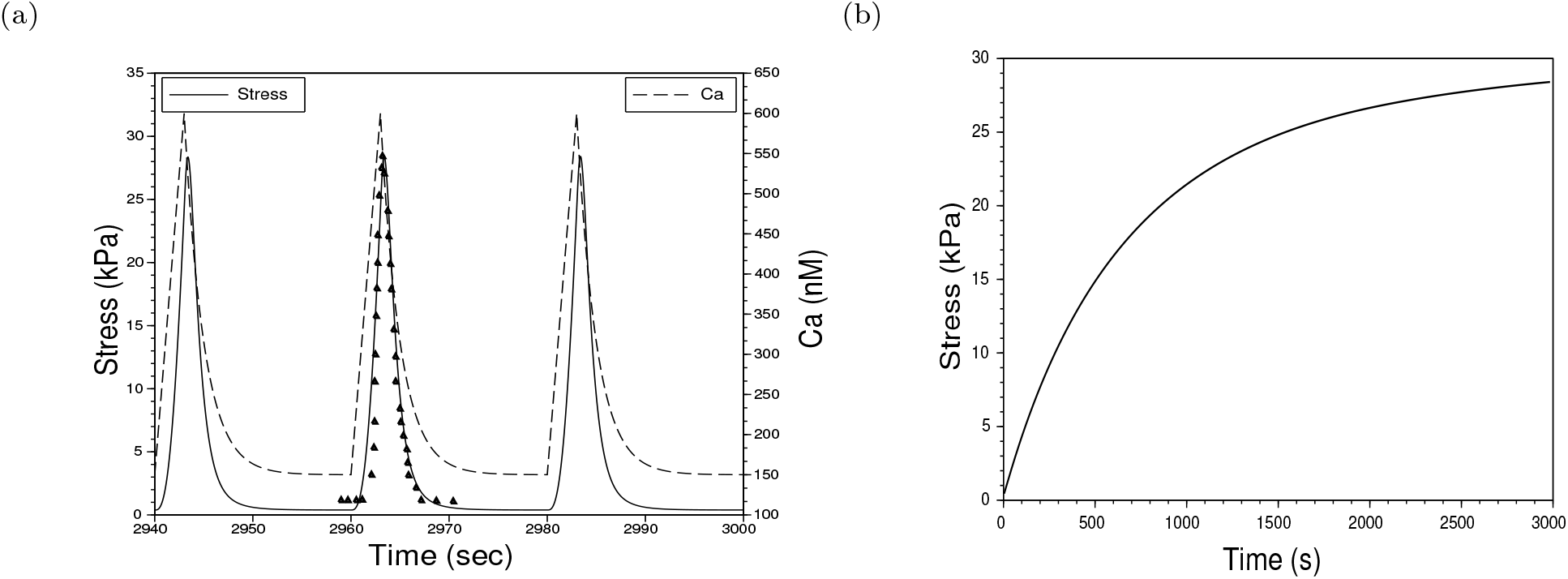
(a) The input [*Ca*^2+^]_*i*_ signal to the model and the model predicted steady-state active stress profile along with the experimental observation (shown with filled triangles) [5, 17]. (b) The evolution of active force with time during the simulation. The solid line as shown in the figure was constructed by picking the peak values of all active force transients during the 3000 seconds of simulation.

**Fig 4:**
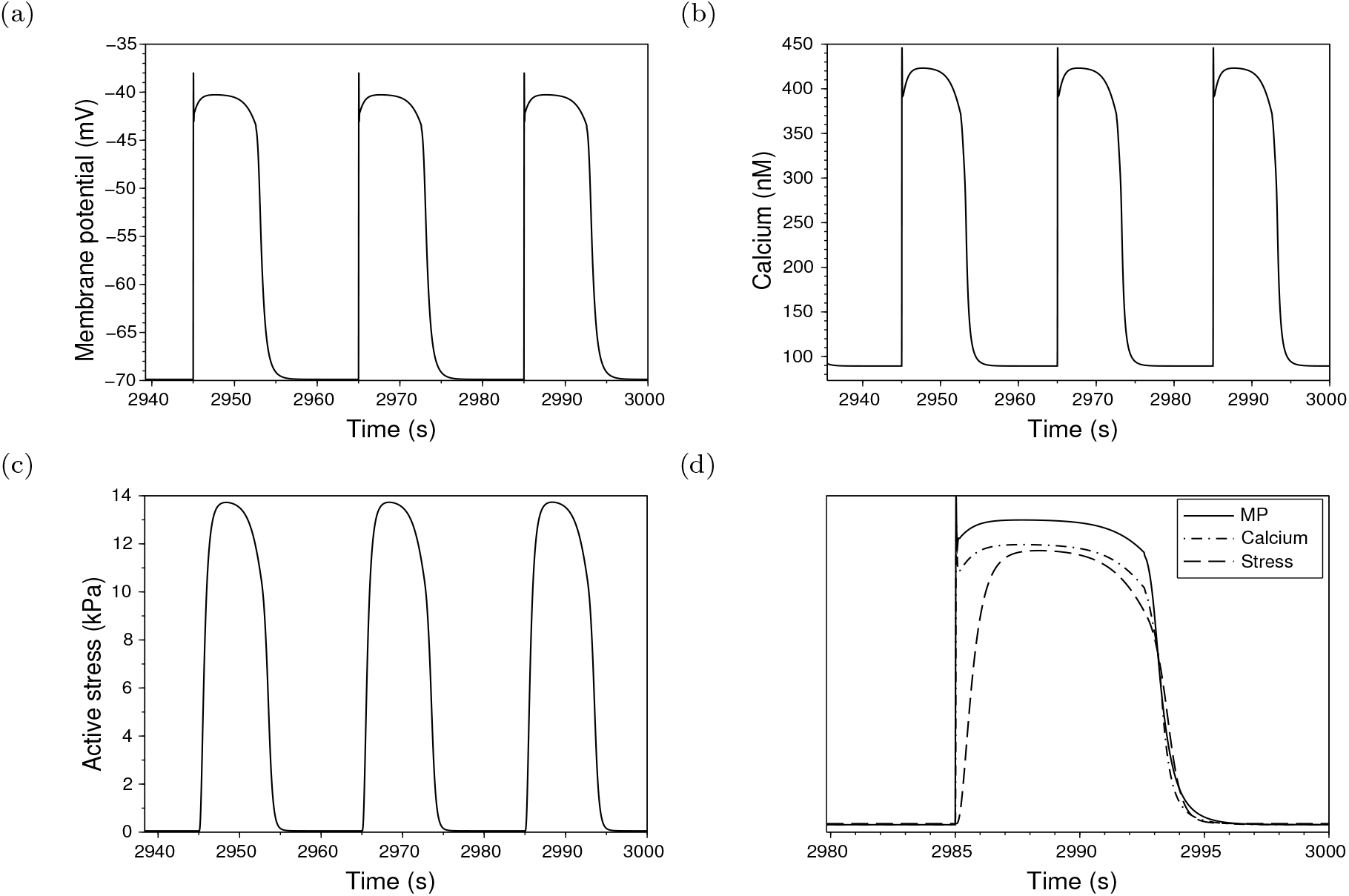
(a) The fluctuation in the MP as a result of the applied electrical pulses. (b) Variation in [*Ca*^2+^]_*i*_. (c) The steady-state active stress profile as a function of time. (d) Comparison of normalised MP, [*Ca*^2+^]_*i*_ and active stress variation in the steady-state.

**Fig 5:**
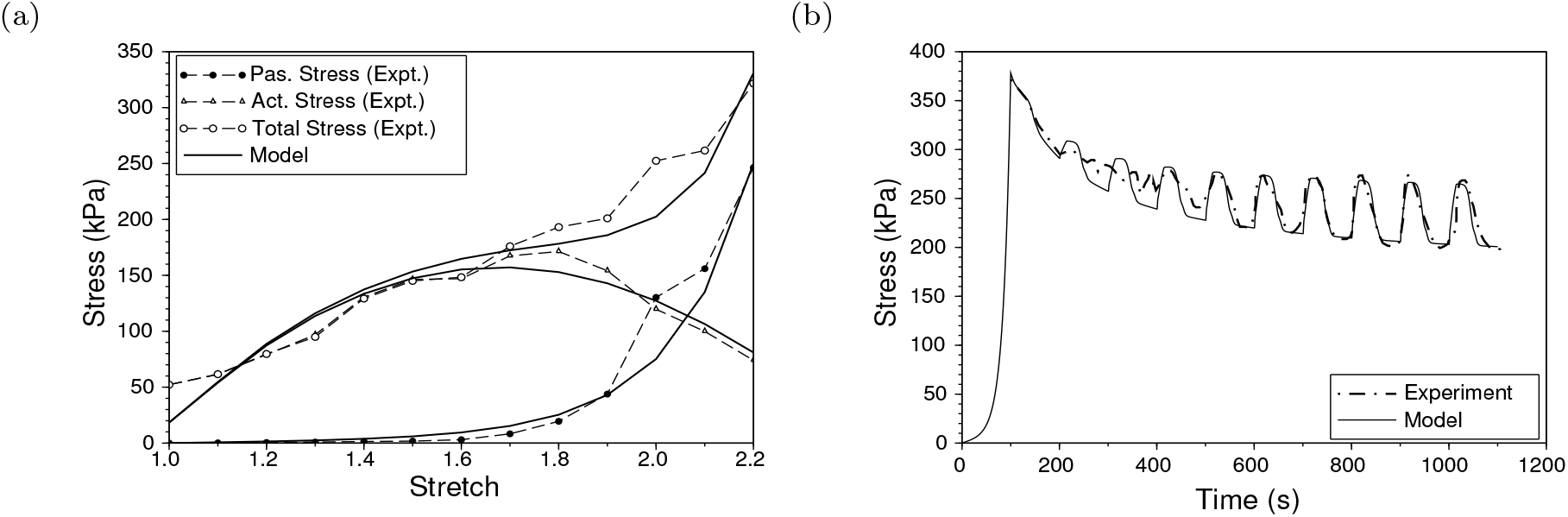
(a) Variation in the active and passive stresses with muscle length [24]. The optimized passive hyperelastic material properties were *c*_1*e*_ = 0.86 kPa and *c*_2*e*_ = 1.43. (b) The active viscoelastic response of a tissue [24]. The tissue stretch was increased up to 2.2 in 100 seconds and then maintained for 1000 seconds to observe stress relaxation. The optimized passive viscoelastic material parameters were *c*_1*e*_ = 0.81 kPa, *c*_1*oe*_ = 1.1 kPa, *c*_2*e*_ = *c*_20*e*_ = 1.43 and *η* = 540 kPa s. The fitting parameter values of the sliding filament model were kept unchanged during both simulations, i.e., *α* = 113.174 kPa, *β* = 0.0353 min^*−*1^ and *κ* = 2098.37 kPa.

Murtada et al. proposed a model to describe this stretch dependency of active force and also validated their work by fitting the model to experimental observations [14, 15, 16]. The primary assumption, for this model, is that the thin actin filaments of a CU with length *L*_*a*_ are longer than the thick myosin filaments (length *L*_*m*_) that gives an overlapping length of *L*_*o*_. The change of length of an activated CU is caused by relative filament sliding (represented by *u*_*fs*_) and the average elastic elongations of the attached cross-bridges (represented by *u*_*e*_). Suppose the length of a CU in its undeformed and deformed states are *L*_*CU*_ and *l*_*CU*_, respectively, then the stretch can be expressed as:

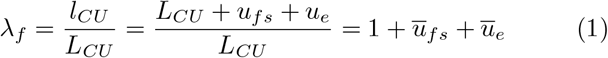

where *ū*_*fs*_ = *u*_*fs*_*/L*_*CU*_ and *ū*_*e*_ = *u*_*e*_*/L*_*CU*_ are taken to be positive for extension. Now, the active stress produced by a CU can be described as:

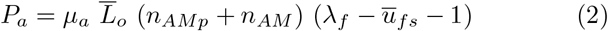

where 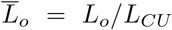 is the normalized overlapping length of actin-myosin filaments, *µ*_*a*_ is a material stiffness parameter and (*n*_*AMp*_ + *n*_*AM*_) is the total number of cross-bridges and latch-bridges inside a CU [16]. The filament overlap, *L*_*o*_ defines the maximum possible number of cross-bridges present in a half CU. Assuming an initial overlap length of *x*_*o*_ between actin-myosin filaments in *Ω*_*o*_, the relative overlap at any instant of time can be expressed as 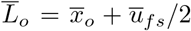 where 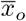 is the normalized initial overlap length between actin-myosin filaments.

The optimum stretch value at which maximum active force is achieved can be expressed as:

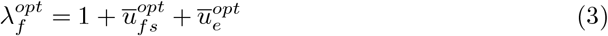

where 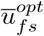 is the optimum filament sliding at which the active force is maximum and 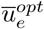 is the corresponding optimum value for the elastic elongation of cross-bridges. Following Murtada et al. [14], a parabolic relation can be formulated for 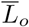 as:

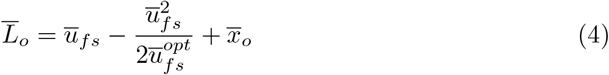

The optimum filament sliding value, 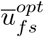, can be calculated using Eq. 3. Now, suppose the steady-state and the optimum active force by the CU are *P*_0_ and *P*_*opt*_, respectively, then the initial overlap of filaments can be calculated as [14]:

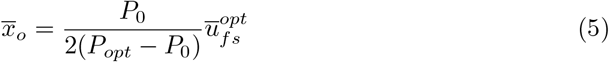

Following Murtada et al. [15], *u*_*fs*_ can be decomposed additively into

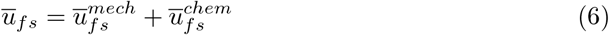

where 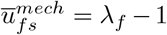 is the filament sliding that arises due to any external mechanical loading or deformation. The filament sliding that is linked to the active cycling cross-bridges is defined through 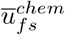. This follows an evolution equation as:

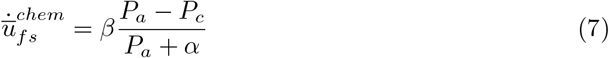

where *P*_*a*_ is the measurable external active stress (Eq. 2) and *P*_*c*_ is the internal stress related to the active cross-bridges which can be defined as:

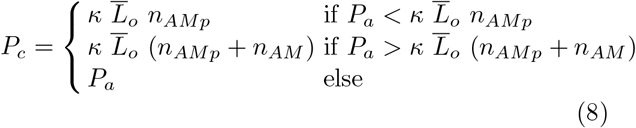

where *α, β* and *κ* are the fitting parameters.

### 2.3 Active finite viscoelastic model

The GI smooth muscle experience a large deformation during normal physiological processes and the tissue stress-strain behaviour is also viscoelastic in nature. To address such complex material behaviour, we previously proposed a finite nonlinear hyper-viscoelastic model [20, 21, 22]. The deformation gradient, for this model, can be decomposed multiplicatively into an elastic and an inelastic part, i.e., **F** = **F**_*e*_**F**_*i*_, and the strain energy can be decomposed into elastic and overstress parts as *ψ* = *ψ*^*E*^+*ψ*^*OE*^. Although it is a three-dimensional model, in one-dimension this model may be thought of as a rheological model analogous to the standard linear solid where both linear springs are replaced with hyper-elastic elements as shown inside the dotted box with a label “passive component” in Fig. 1a. The decomposed parts of the total strain energy, *ψ*^*E*^ and *ψ*^*OE*^, are associated with the parallel and in-series hyperelastic elements, respectively. Now, assigning Humphrey strain energy functions to 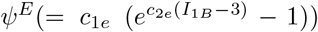 and 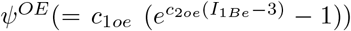, the second PK stress tensor for this model can be derived as:

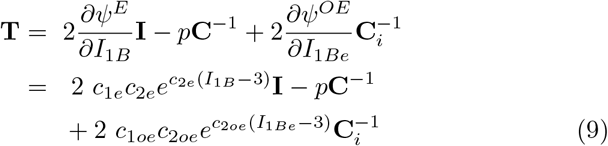

where *c*_1*e*_, *c*_1*oe*_, *c*_2*e*_, and *c*_2*oe*_ are material stiffness parameters, (*I*_1*B*_, *I*_2*B*_) and (*I*_1*Be*_, *I*_2*Be*_) are the first and second invariants of **B** and 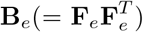, respectively, *p* is a pressure term that is used to enforce the incompressibility constraint, and 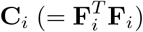 is the inelastic component of **C** [20].

For a purely mechanical system, the second law of thermodynamics takes the form [8]:

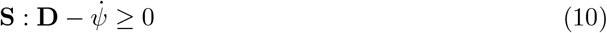

where **D** is the symmetric part of **L**. Following the steps outlined by Panda and Buist [20], Eq. 10 can be reduced to

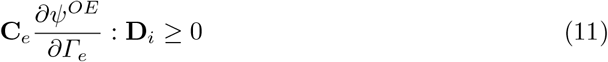

where *Γ*_*e*_ is the elastic part of strain tensor defined in an intermediate configuration, *Ω*_*i*_, that exist in between the reference and current configuration during the finite deformation of the continuum body. One of the simplest ways to satisfy Eq. 11 is to express **D**_*i*_ as:

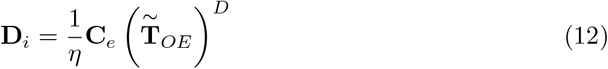

where 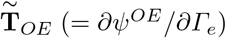 is the overstress part of second PK stress tensor defined in *Ω*_*i*_ and *η* is the viscosity of the material. Now, using Eq. 12 and the relation 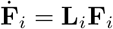, an evolution equation for **C**_*i*_ can be derived as:

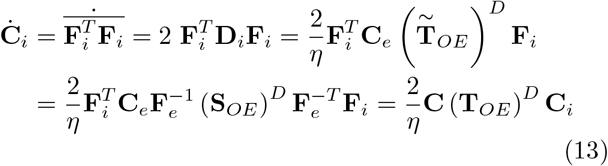

We further extended this model by adding an extra parallel arm for active tissue mechanics as shown in Fig. 1a. We denote this model as the active finite nonlinear hyper-viscoelastic model (AFNHVM). Here, we assume that the CU, surrounded by a fibrous matrix, generates active contractile forces, whereas the fibrous matrix imparts passive forces that resist deformation. Thus, the complete AFNHVM model can be thought as a passive viscoelastic component in parallel with an active component as shown schematically in Fig. 1a. The contractile element (see Fig. 1a) incorporates the actin-myosin interaction model and sliding filament model in it and provides the active force generated in the SMC as an output.

The actin-myosin interaction model, given an [*Ca*^2+^]_*i*_ fluctuation, calculates the number of cross-bridges, i.e., (*n*_*AMp*_ + *n*_*AM*_), inside a SMC. This is a key input to the sliding filament model. The other input needed by the sliding filament model is the stretch in the direction of the CU, i.e., *λ*_*f*_. Suppose, the alignment direction a CU inside a SMC is given by an unit vector **f**, then the stretch in the direction of the CU can be computed as:

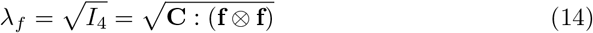

where *I*_4_ = **C** : (**f** ⊗ **f**) is the pseudo-invariant of the deformation tensor, **C** is the deformation tensor experienced by the SMC and ⊗ is the dyadic product. With these key inputs, the sliding filament model calculates the active force which can then be converted into an active first PK stress tensor as **P**_*active*_ = *P*_*a*_.(**f** ⊗ **f**) where *P*_*a*_ is the active force produced by the sliding filament model. Now, this active stress tensor can be added to the passive stress tensor of the nonlinear viscoelastic model to obtain total stress. Using Eq. 2 and Eq. 9, the total first PK stress tensor can be written as:

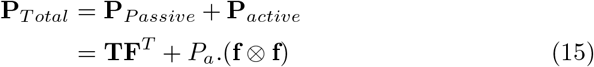

This completes the coupling between different models to form the AFNHVM. However, an cellular electrophysiology model is needed to compute the fluctuation in [*Ca*^2+^]_*i*_ which is a key input to the actin-myosin interaction model.

### 2.4 Smooth muscle cell model

The biophysical based gastric SMC model proposed by Corrias and Buist was utilized in this work [1]. Although other SMC model exists in the literature, they are either inconsistent with current experimental data or phenomenological, with model parameters of no biological significance [11, 25]. The Corrias and Buist SMC model is based on the Hodgkin-Huxley-like description of the cell membrane with eight ion channels conductances and has been validated against canine gastric SMC data. This model, given an electrical stimulus as an input, calculates the fluctuation in membrane potential (MP) as well as in [*Ca*^2+^]_*i*_. The interested reader is referred to the original work for further details [1].

We coupled this electrophysiology model with the AFNHVM model to formulate an electromechanical framework which can be used to calculate the total stress and strain in a SMC provided an input electrical stimulus. The complete coupling process for the AFNHVM is depicted in Fig. 1b.

## 3 Results

To demonstrate the efficacy of the AFNHVM model, it was utilized to recreate experimental results performed of gastric tissue strips under different loading conditions.

Gerthoffer et al. performed experiments on canine tissue strips obtained from the circular muscle layer of the proximal colon to measure [*Ca*^2+^]_*i*_, myosin phosphorylation and active force values simultaneously during tissue contraction induced using a *K*^+^-PSS solution [6]. Their experimental observations are shown in Fig. 2b and 2c. During the experiment, [*Ca*^2+^]_*i*_ was found to increase rapidly to a peak value and then remained constant with time. To recreate this experimental observation, we provided a step variation in [*Ca*^2+^]_*i*_ as an input to the AFNHVM model (see Fig. 2a) and then optimized the model parameters to best-fit the active stress profile. The peak value of [*Ca*^2+^]_*i*_ was set in such a way that the model predicted myosin phosphorylation value matched the experimental observation (see Fig. 2b). The parameters of the sliding filament model, except *α, β* and *κ*, require an active force-length relationship for their identification. As Gerthoffer et al. did not report any active force-length relationship in their work, we adopted the experimental results of Price et al. to estimate these parameters. In their study, Price et al. dissected taenia coli strips from the cecum and measured the active force value produced by the tissue at different stretching lengths during spontaneous contraction (see Fig. 2d) [24]. From this experimental data, 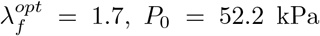 and *P*_*opt*_ = 171.3 kPa can be extracted. We set the value of 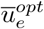 at 0.02 [14]. Using Eq. 3 and Eq. 5, 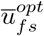 and 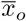 were calculated to be 0.68 and 0.149, respectively. Now, using Eq. 2 and Eq. 3, the parameter *µ*_*a*_ can be computed as 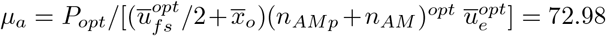 MPa where the myosin phosphorylation level at the optimum length was set at (*n*_*AMp*_ + *n*_*AM*_)^*opt*^ = 0.24 (see Fig: 2b). Finally, the fitting parameters of the sliding filament model, *α, β* and *κ*, were optimized by minimizing the distances between the experimental data points (see Fig. 2c) and simulation results using Nelder-Mead algorithm in Unfit (Release 3.0, Computational Bioengineering Lab, NUS). The model prediction for the evolution of active stress with respect to time and the active stress-length relationship are depicted in Fig. 2c and 2d, respectively. The complete active response for different muscle lengths (*λ*_*f*_ = 1.0 *−* 2.2) from the time of activation up to 200 seconds are presented in Fig. 2e and 2f.

Experiments were performed by Ozaki et al. to measure [*Ca*^2+^]_*i*_ and active force values simultaneously during spontaneous contraction of SMC excised from a canine stomach [17, 18]. Fig. 3a illustrates the variation of active stress with [*Ca*^2+^]_*i*_ in a tissue strip. The active force value reported by the authors were in the unit of force (mN). Thus, we converted that into the stress units by dividing by the specimen area. We assumed the thickness of the circular muscle layer to be 0.5 mm and then multiplied that with the reported specimen width to get the area [6, 27]. Furthermore, the [*Ca*^2+^]_*i*_ data was presented in terms of fluorescence ratio and that was converted into a suitable unit (nM) following the procedure as outlined by Gajendiran and Buist [5]. A train of [*Ca*^2+^]_*i*_ transients was generated to simulate the presence of slow wave activity (see Fig. 3a) and then furnished to the model as an input. The resulting active stress generated by the model was found to evolve with time and achieved an equilibrium value after 3000 seconds of simulation (see Fig. 3b). The steady-state active force value along with the experimental observation are presented in Fig. 3a.

Next, we assigned a train of transient electrical pulses to the model as an input and monitored the corresponding changes in MP, [*Ca*^2+^]_*i*_, and active stress (Fig. 4). The electrical stimulus, representing an slow wave that depolarises SMC, was constructed following Corrias and Buist (see supplementary material) [1]. Slow waves occur at a frequency of 3 cycles per minute in stomach [1, 2]. Thus, for this simulation, the stimulus frequency was set to the same. Fig. 4a illustrates the resulting fluctuation in the MP which consists of a rapid upstroke to a peak value of -38 mV, followed by a prolonged plateau phase around -40 mV, and a resting membrane potential of -70 mV. These MP values are in close agreements with the experimental observations (experimental peak, plateau and resting membrane potentials are -34 mV, -42 mV and -71 mV, respectively [18]). The [*Ca*^2+^]_*I*_ variation followed the MP closely as shown in Fig. 4b. A variation of 300 nM was observed in [*Ca*^2+^]_*i*_ from the resting level to the peak level which agrees with the available experimental data [1]. The active stress was found to evolve with time and achieved a steady-state value after 3000 seconds of simulation (see supplementary material). The transient active force profile was smooth without any rapid upstroke (see Fig. 4c) and it reached the peak value after the maximum increase in [*Ca*^2+^]_*i*_ was achieved (see Fig. 4d). Experimental data suggests that the active force varies in between 0.6-0.9 gram during a spontaneous contraction of SMC [18]. According to Zhao et al. [27], the muscle layer thickness of pig intestine is 0.9 ± 0.06 mm, and as per Liao et al. [12], the muscle layer thickness is 1.03 ± 0.08 mm. Assuming the circular and longitudinal muscle layers are of the same thickness, the circular muscle layer thickness was calculated to be in between 0.42 mm to 0.55 mm. With these extreme values for the thickness, the active stress values can be calculated to be in between 14.01-21.02 kPa and 10.7-16.05 kPa, respectively. Our predicted active stress was 13.51 kPa which lies with in the observed range. A plot consolidating the normalized variations in the MP, [*Ca*^2+^]_*i*_ and active stress is given Fig. 4d for comparison which agrees well with the experimental observation of Ozaki et al. (see Fig.1 of [18]).

Smooth muscle passive stress-stretch relations are highly nonlinear and the stress increases exponentially with stretch [20]. Fig. 5a shows the variation in the passive and active stress with stretch in a porcine taenia coli [24]. It is evident from the figure that the passive stress is nonlinear and the active stress follows a bell-shaped relationship with stretch. In addition, the active stress curve is almost symmetrical about the optimal point and the stress decreases in either direction form 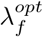. In the absence of any time-dependent experimental data, we considered a hyperelastic material for this tissue. Hyperelastic material responses can be obtained form Eq. 9 by setting the values of the parameters *c*_1*oe*_ and *c*_2*oe*_ equal to zero. The elastic stiffness parameters, i.e., *c*_1*e*_ and *c*_2*e*_, were optimized first to obtain the nonlinear passive tissue properties and then a train of electrical pulses was feed to the model to obtain the active responses (see supplementary material for the pulse profile). The resulting active tissue behaviour is shown in Fig. 5a. For this simulation, the applied stretch was increased in steps of 0.1 and then kept constant for 200 seconds in each step. During this time, electrical pulses were applied continuously to obtain the steady-state behaviour of active stress. Finally, the total stress was calculated by adding the active and passive stress values together.

An experimental study of Price et al. revealed the passive viscoelastic behaviour along with the superimposed active behaviour of a porcine taenia coli during spontaneous contractions [24]. In their experiment, the authors stretched a tissue strip to a higher length in 100 seconds and then maintained that stretch for 1000 seconds to observe changes in the passive as well as in the active tissue behaviour. During the constant stretching phase, the passive stress was found to relax from an instantaneous value towards an equilibrium value (see Fig. 5b). On the other hand, the active stress gradually evolved with time and built up to a steady state value. We optimized the material parameters of Eq. 9 to obtain the passive viscoelastic properties. The experimental data suggests that each cyclic contraction is approximately 100 seconds long (see Fig. 5b). Thus, we constructed an electrical pulse of the same length and adjusted its magnitude and profile in such a way that the corresponding contractile rhythms would match the experimental observation (see supplementary material for the stimulus profile). The model response for this simulation is shown in Fig. 5b which agrees well with the experimental data.

Systemic sclerosis (SS) is a connective tissue disease in which the GI tract often gets affected. Pedersen et al. examined the in-vivo healthy and SS diseased tissue properties of the human intestine and showed that in SS the intestine causes dilation with impaired muscle function [23]. In their study, impedance planimetry and balloon distension were used to obtain the active and passive tension-length curves. Thus, we considered a hyperelastic cylinder inflation protocol to recreate this experimental observation. A brief discussion about the theory for a cylinder inflation with active stress and the simulation set up is given in the supplementary material. We considered a cylinder with inner radius of 7.31 mm to represent the intestine and then inflated that cylinder to make the inner radius 17.67 mm. These radii were calculated using the cross-sectional data as reported by the authors during inflation. A train of transient electrical pulses was applied during inflation to elicit the active tissue behaviour and then the corresponding radial stress at the inner wall of the cylinder was monitored. The model predicted active and passive radial stresses at the inner wall of the cylinder during inflation are presented in Fig. 6a along with the experimental observations. For diseased tissue simulation, the inner radius of the cylinder was calculated to be 6.85 mm which was then inflated up to 19 mm. The corresponding diseased tissue behaviour is depicted in Fig. 6b. These results clearly indicate that both active and passive tissue properties change during SS.

**Fig 6:**
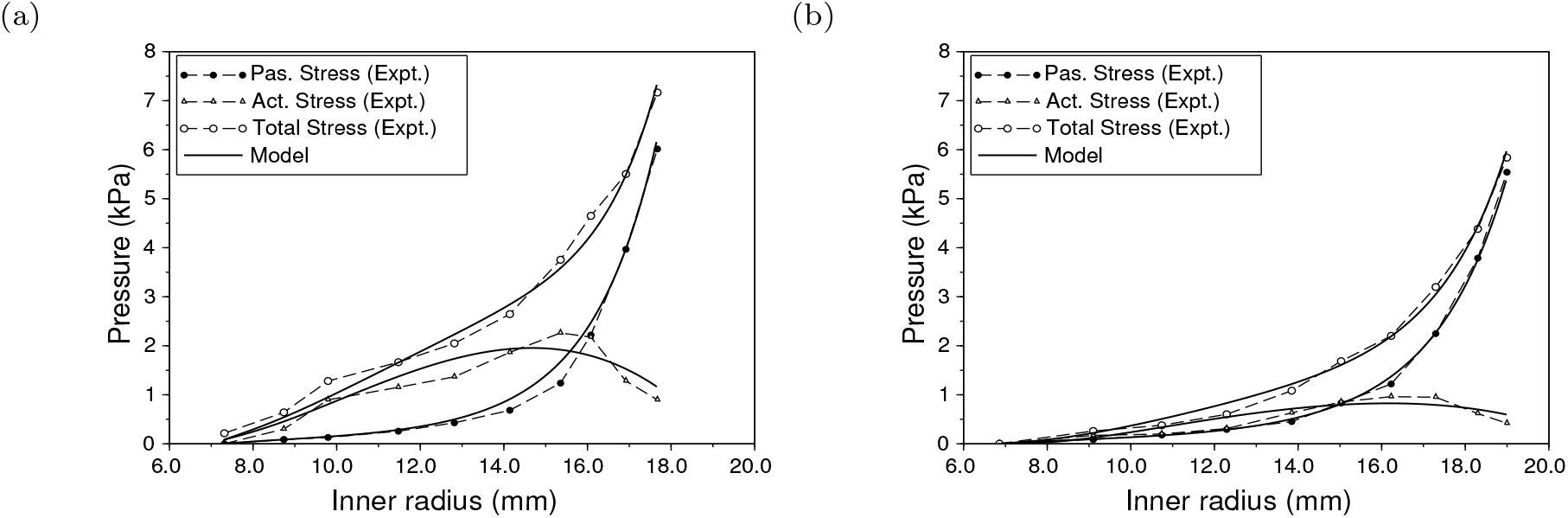
The in-vivo active and passive tissue properties of human intestine in health (a) and in disease (b). The dotted lines are the experimental observation, whereas the solid lines depict model responses. The optimized passive viscoelastic properties for the healthy tissue were *c*_1*e*_ = 0.6 Kpa and *c*_2*e*_ = 1.05. The corresponding values for the diseased tissue were 0.6 kPa and 0.75, respectively. The fitting parameters for the sliding filament model were kept unchanged both for healthy and diseased tissue simulations, i.e., *α* = 113.174 kPa, *β* = 0.0353 min^*−*1^ and *κ* = 2098.37 kPa. For healthy tissue simulation, the active tension-length depended parameters were set at 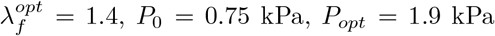, *P*_0_ = 0.75 kPa, *P*_*opt*_ = 1.9 kPa and (*n*_*AMp*_ + *n*_*AM*_)^*opt*^ = 0.24. Similarly, for the diseased tissue simulation, they were 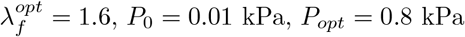 and (*n*_*AMp*_ + *n*_*AM*_)^*opt*^ = 0.24.

**Fig 7:**
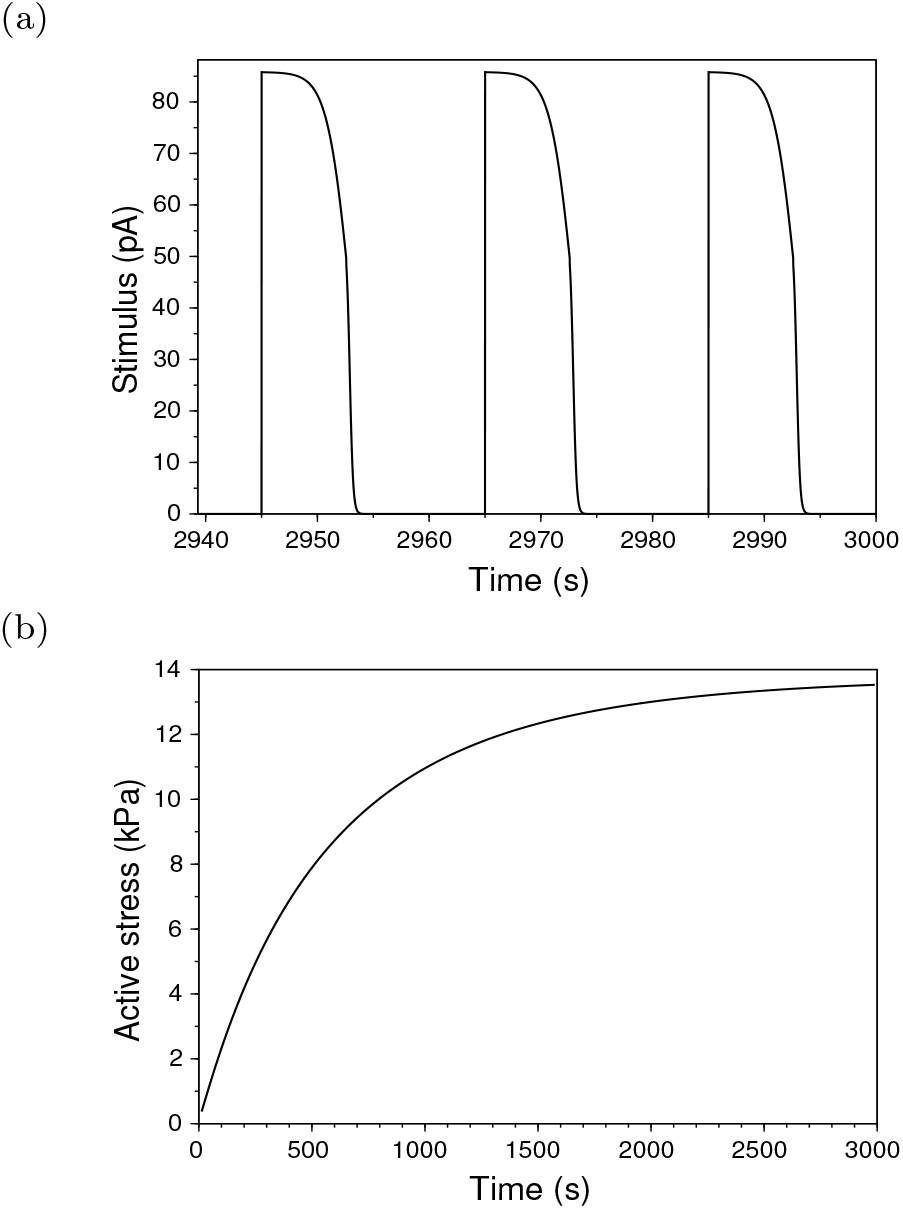
(a) A train of electrical pulses that served as an input to the model. (b) Evolution of active force with time.

**Fig 8:**
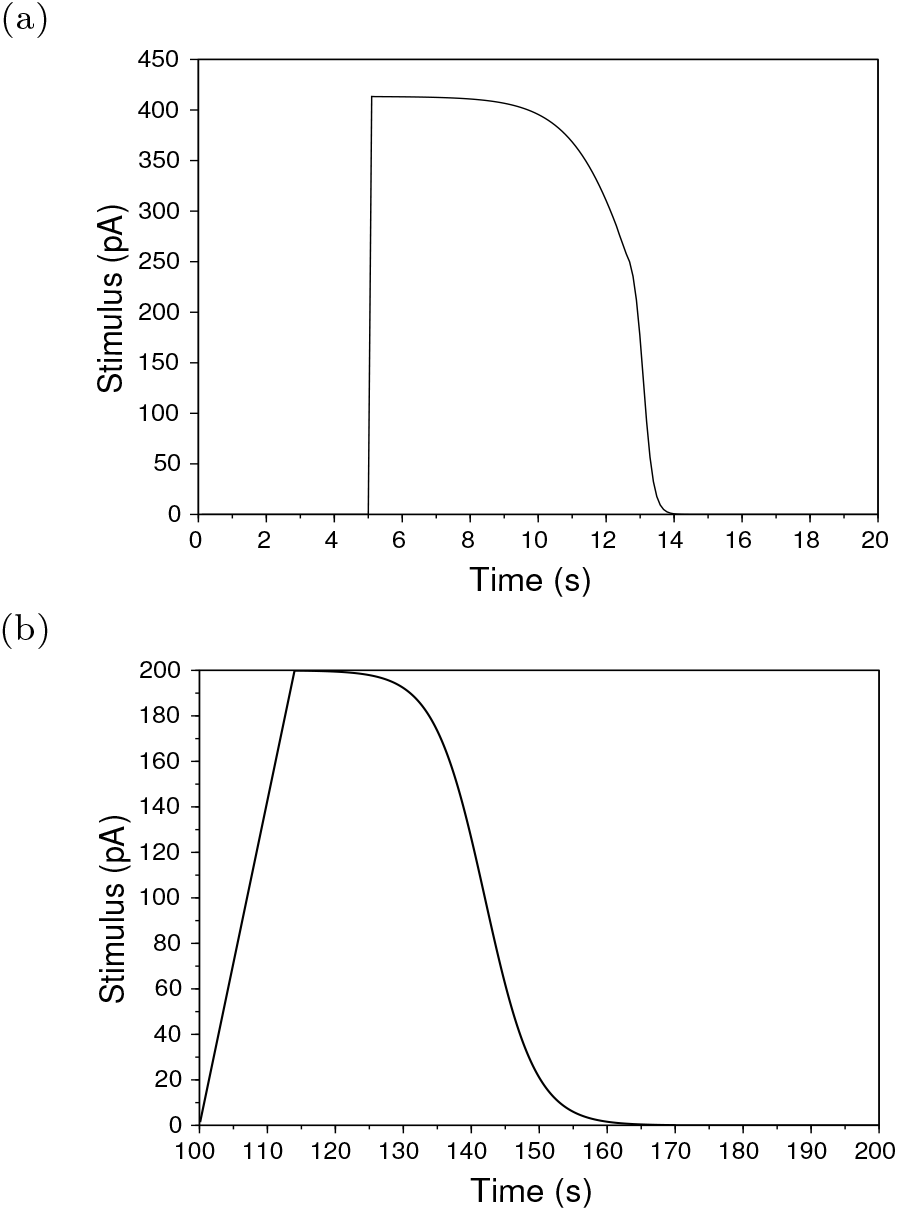
(a) The input electrical pulse used for active stress generation. (c) The input electrical signal used during stress relaxation simulation.

## 4 Discussion and conclusions

This study presents an electromechanical model demonstrating the transduction process of cellular electrical activity into mechanical deformation. The ability to link cellular activities to motility provides an opportunity to study complex subcellular mechanisms and would also be helpful for mechanical studies where the GI luminal content movement is modelled by imposing mechanical deformation to the wall from imaging evidence [19]. A coupled electromechanical model would also be advantageous for studying the effects of channelopathies on motility.

The SMC cannot generate electrical activity on their own and, thus, they depend on external stimuli. The Interstitial cell of Cajal (ICC) are the natural pacemaker of the GI wall and are responsible for the omnipresent electrical activity intrinsic to the GI wall, whereas the enteric nervous system provides an additional extrinsic level of control. Although a biophysical model exists in the literature for ICC [2], that was not considered in this study as the ICC are not the sole contributor to SMC depolarization. Therefore, a train of electrical pulses was used as an input in our study. However, an ICC model can easily be coupled with the existing model via a gap junction conductance.

In the experimental study of Ozaki et al., the muscle tension decreased more rapidly than the [*Ca*^2+^]_*i*_ which signifies the existence of some inactivation mechanism in the CU [17]. Some studies suggest that the activation of the enzyme myosin light chain phosphatase and calcium desensitization may contribute to this rapid relaxation of contractile force as compared to the [*Ca*^2+^]_*i*_ decline [17, 5]. As these inactivation mechanisms were not considered in our study, the results, as in Fig. 4d, are not consistent with those aspects of the experimental observations. Extending the current model to incorporate these additional regulatory pathways would further enhance the model.

We modified the active force model of Murtada et al. to reduce the number of fitting parameters and then estimated the parameter values using different experimental data sets. Different experimental observations, e.g., contractile forces in canine stomach tissues and in porcine colonic tissues, were recreated using the same model without altering any fitting parameters and simply by changing the magnitude and profile of the input stimulus. This signifies the effectiveness and versatility of the proposed method to model GI electromechanics. A sample implementation of the model is available from the authors upon request.

## Acknowledgements

We would like to thank Dr. Alberto Corrias for his valuable comments. We also gratefully acknowledge the President’s Graduate Fellowship, NUS for funding support.

## A Cylinder inflation with active stress

Consider a cylinder with an internal radius *A*, external radius *B* and length *L*. Suppose, this cylinder has been inflated with an internal pressure *P*_*i*_, and after inflation, the internal, external radii and length are *a, b* and *l*, respectively. Now, the continuum body in its inflated configuration should satisfy the equilibrium equation. The equilibrium equation for a symmetric circular geometry in the cylindrical coordinate system is given as:

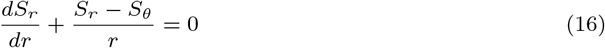

where *S*_*r*_ and *S*_*θ*_ are the components of the Cauchy stress in the radial and circumferential direction, respectively. The quantity *r* represents the radial coordinate of any material point in the inflated configuration. Considering an incompressible material, the deformed and undeformed radial coordinates (i.e., *r* and *R*) can be related to each other as:

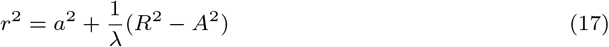

where *λ* (= *l/L*) is the axial stretch. Now, the stretches in the cylindrical coordinate system can be expressed as:

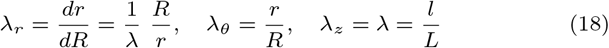

These are also the principal stretches and satisfy the incompressibility condition, i.e., *λ*_*r*_*λ*_*θ*_*λ*_*z*_ = 1.

Let’s consider a Humphrey’s material for the cylinder.The Cauchy stress for this material can be written as:

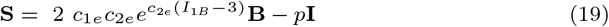

where *p* is the hydrostatic pressure necessary to satisfy the incompressibility condition.

Eq. 16 is a boundary value problem. It can be solved using Eq. 19 with specified boundary conditions at the internal and external surfaces of the cylinder. Suppose, the internal surface of the cylinder is subjected to an internal pressure *P*_*i*_, whereas the external surface is stress-free. Then, Eq. 16 transforms into:

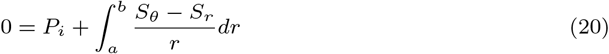

This is a nonlinear equation which can be solved numerically to obtain the internal pressure in the inflated configuration. Now, the Cauchy stresses across the wall of the cylinder in the inflated configuration can be expressed as:

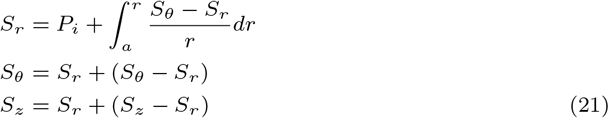

The active stress value, as in Eq. 2, with an appropriate transformation can be added to the passive stress value, as in Eq. 19, to obtain the combined effect. For the inflation testing simulation, as shown in Fig. 6, the muscle fibers were assumed to be aligned along the circumferential direction of the cylinder. Thus, before solving the equilibrium equation, the active stress value was added to the circumferential component of Cauchy stress.

## Supplementary material

### Simulation environment for different tests

The input electrical pulses, used in the simulation as described in Fig. 4, to elicit active stress behaviour is given in Fig. 7a. The GI slow waves occur at a frequency of 3 cycles per minute in stomach. Thus, for this simulation, the stimulus frequency was set to three. Fig. 7b shows the evolution of active force as a function of time. The solid line as shown in figure was constructed by picking the peak values of all active force transients (see Fig. 4c) during the 3000 seconds of simulation.

Fig. 8a shows the input electrical stimulus used in the simulation, as in Fig. 5a, to generate active forces and Fig. 8b is the one used during stress relaxation simulation as in Fig. 5b.

